# The X-linked intellectual disability gene product and E3 ubiquitin ligase KLHL15 degrades doublecortin proteins to constrain neuronal dendritogenesis

**DOI:** 10.1101/2020.10.02.324285

**Authors:** Jianing Song, Ronald A. Merrill, Andrew Y. Usachev, Stefan Strack

**Affiliations:** Department of Neuroscience and Pharmacology and the Iowa Neuroscience Institute, University of Iowa, Iowa City, Iowa 52242

**Keywords:** Kelch-like 15, KLHL15, E3 ubiquitin ligase, protein phosphatase 2A, signal transduction, doublecortin, doublecortin-like kinases, microtubule-associated protein, ubiquitination, proteasomal degradation, protein turnover, neurite outgrowth, dendritic complexity, pulse-chase, HaloTag

## Abstract

Proper brain development and function requires finely controlled mechanisms for protein turnover and disruption of genes involved in proteostasis is a common cause of neurodevelopmental disorders. Kelch-like 15 (KLHL15) is a substrate adaptor for cullin3 (Cul3)-containing E3 ubiquitin ligases and *KLHL15* gene mutations were recently described as a cause of severe X-linked intellectual disability. Here, we used a bioinformatics approach to identify a family of neuronal microtubule-associated proteins (MAPs) as KLHL15 substrates, which are themselves critical for early brain development. We biochemically validated doublecortin (DCX), also an X-linked disease gene, and doublecortin-like kinases 1 and 2 (DCLK1/2) as *bona fide* KLHL15 interactors and mapped KLHL15 interaction regions to their tandem DCX domains. Shared with two previously identified KLHL15 substrates, a FRY tripeptide at the C-terminal edge of the second DCX domain is necessary for KLHL15-mediated ubiquitination of DCX and DCLK1/2 and subsequent proteasomal degradation. Conversely, silencing endogenous KLHL15 markedly stabilizes these DCX domain-containing proteins and prolongs their half-life. Functionally, overexpression of KLHL15 in the presence of wild-type DCX reduces dendritic complexity of cultured hippocampal neurons, whereas neurons expressing FRY-mutant DCX are resistant to KLHL15. Collectively, our findings highlight the critical importance of the E3 ubiquitin ligase adaptor KLHL15 in proteostasis of neuronal MAPs and identify a regulatory network important for development of the mammalian nervous system.

## INTRODUCTION

Ubiquitination is a highly coordinated multi-step enzymatic cascade that requires the concerted action of three key factors: E1 ubiquitin-activating enzymes, E2 ubiquitin-conjugating enzymes, and E3 ubiquitin ligases (1), many of which are strongly implicated the pathogenesis of neurodevelopmental disorders (2-4). E3 ubiquitin ligases have received particular attention, as they confer substrate specificity of the ubiquitination reaction by catalyzing the transfer of ubiquitin to proteins destined for degradation or other ubiquitin-dependent fates (5-7).

Recent studies have identified *KLHL15* as a X-linked intellectual disability (XLID) gene. Inactivating *KLHL15* mutations are associated with various brain abnormalities as well as developmental and behavioral disorders in male patients (8-11). Localized on chromosome Xp22.11, *KLHL15* encodes a member of the Kelch-like (KLHL) family of proteins that function as adaptors for Cullin3 (Cul3)-based E3 ubiquitin ligases to target specific substrates to the ubiquitin-proteasome system (12). Numbering more than 40 in humans, KLHL proteins exert a wide range of biological functions, while genetic mutations and abnormal expression of *KLHL* genes have been linked to diverse diseases, ranging from cardiovascular disorders to cancer (12-14).

We have previously demonstrated that KLHL15 mediates the ubiquitination and subsequent proteasomal degradation of the B*’β* (B56*β*, PR61*β*, PPP2R5B) regulatory subunit of the protein Ser/Thr phosphatase 2A (PP2A). KLHL15-mediated B*’β* degradation facilitates the formation of alternative PP2A holoenzymes by promoting the exchange with more than a dozen other regulatory subunits (15). PP2A/B*’β* is highly enriched in the mammalian nervous system (16-18) and plays key roles in striatal dopamine biosynthesis (16), hippocampal long-term potentiation (19), and multiple intracellular signaling pathways in response to growth factor stimulation (20-22). We also identified a tyrosine residue (Y52) within a phylogenetically well-conserved tripeptide motif (FRY) in vertebrate B*’β* as being necessary for KLHL15 binding and KLHL15-mediated B*’β* downregulation (15). Since then, work by others identified a critical FRY tripeptide in the degron of a second KLHL15 substrate, the DNA repair protein CtIP/RBBP8 (23).

Because of *KLHL15*’s link to severe intellectual disability and its ranking as one of the most clinically significant X-linked disease genes (8,24), we set out to uncover novel neuronal substrates of the E3 ubiquitin ligase by primary and secondary structure searches. This screen and subsequent biochemical validation identified doublecortin (DCX) and two doublecortin-like kinases (DCLK1, DCLK2), members of a family of neuronal microtubule-associated proteins (MAPs) that share a characteristic domain structure of N-terminal tandem DCX domains. These tandem DCX domains stabilize microtubule by enhancing tubulin polymerization, inhibiting tubulin catastrophe, and promoting microtubule bundling (25-28). DCX plays essential roles in neurogenesis, neuronal migration, axonal/dendritic wiring, and cargo transport therefore ensuring normal brain development (28,29). The *DCX* gene is located on the X chromosome and *DCX* mutations cause classic lissencephaly (agyria) in males and subcortical band heterotopia (also called double cortex syndrome) primarily in females as a result of neuronal migration defects (30). Both disorders typically manifest with intellectual disability, epilepsy, language impairment, and global developmental delay (30). DCLK1 and DCLK2 feature a C-terminal Ser/Thr kinase domain and play both common and distinct roles in neurodevelopment. Both kinases associate with microtubules and stimulate microtubule polymerization and dendritic development independent of their kinase activity (31-33). DCLK1 and DCX synergistically regulate neuronal migration, axonal outgrowth, and hippocampal development (34-36), whereas DCLK2 works in concert with DCX to organize hippocampal lamination and prevent spontaneous seizures (37).

This report documents that three members of the DCX family of neuronal MAPs, are subject to KLHL15-mediated downregulation. KLHL15-dependent polyubiquitination and proteasomal degradation requires a degron containing both N-terminal DCX domains and a conserved FRY tripeptide. Functionally, we show that KLHL15 expression reduces dendritic arborization in primary hippocampal cultures unless neurons also express DCX with a stabilizing FRY mutation. Our study uncovers a novel role of the E3 ubiquitin ligase adaptor KLHL15 as a negative regulator of DCX protein abundance and dendritogenesis and suggests possible mechanistic connections in X-linked neurodevelopmental disorders.

## RESULTS

### KLHL15 regulates signal transduction in hippocampal neurons by PP2A-dependent and independent mechanisms

To determine whether KLHL15 possesses functions in neurons independent of B*’β*, a neuron-enriched regulatory subunit of PP2A (16-18), we investigated the effect of KLHL15 on activation of extracellular signal-regulated kinases (ERKs). We previously reported that the B*’β*-containing PP2A holoenzyme potentiates nerve growth factor (NGF) signaling at the tropomyosin-related kinase A (TrkA) receptor level to sustain ERK signaling and neuronal differentiation of neuroendocrine PC12 cells (22). To address the hypothesis that PP2A/B*’β* may similarly potentiate TrkB receptor activity in central neurons, we measured ERK activity in transiently transfected primary hippocampal neurons from E18 rat embryos using a dual luciferase reporter assay (PathDetect Elk1 reporter, Fig. 1A) (22). In GFP control-transfected neurons, a 3 h incubation with a sub-saturating concentration of brain-derived neurotrophic factor (BDNF) resulted in a ∼17-fold increase in Elk1 transcriptional activity (Fig. 1B). Compared to control, overexpression of GFP-KLHL15 attenuated ERK activation by 2.5-fold, whereas B*’β* overexpression amplified the BDNF response by 3-fold (Fig. 1B). Conversely, silencing of endogenous KLHL15 amplified BDNF signaling to ERK by 1.5-fold over control shRNA-transfected cells, while shRNA targeting endogenous B*’β* nearly eliminated the BDNF response (Fig. 1C). These results suggest that KLHL15 inhibits neurotrophin signaling in hippocampal neurons primarily by targeting PP2A/B*’β* for proteasomal degradation. The neuromodulator pituitary adenylate cyclase-activating polypeptide (PACAP) signals through the type 1 G protein–coupled receptor PAC1 to ERK via cAMP–dependent activation of Rap1 (38) (Fig. 1A). In control-transfected neurons, PACAP at sub-saturating concentrations induced Elk1 transcriptional activity by ∼5-fold. Overexpression of GFP-KLHL15 dampened this response by ∼2-fold, whereas KLHL15 silencing amplified PACAP signaling to ERK by ∼3-fold. In contrast to BDNF stimulation, neither overexpression nor knockdown of PP2A/B*’β* significantly affected the reporter response to PACAP (Fig. 1, D and E), suggesting the involvement of a distinct KLHL15 substrate.

**Figure 1.**
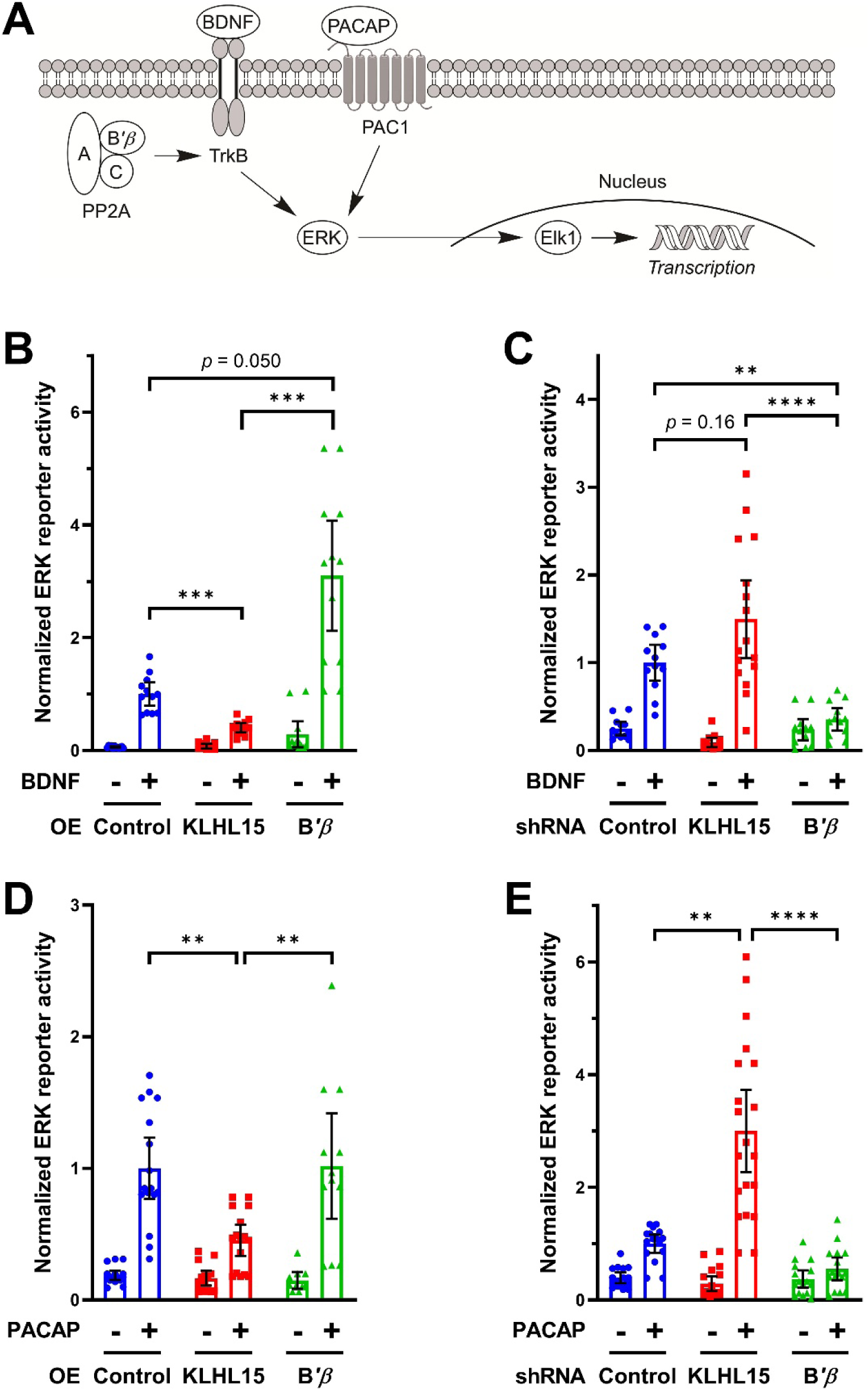
KLHL15 modulates BDNF- and PACAP-mediated ERK activation in hippocampal neurons by different mechanisms. *A*, schematic of BDNF and PACAP signaling resulting in Elk1-mediated luciferase activity as a reporter of ERK activation. *B–D*, hippocampal cultures from E18 rat embryos transfected with the indicated cDNAs or shRNAs together with dual-luciferase reporter plasmids (PathDetect Elk1 reporter) were stimulated (+) with BDNF (25 ng/ml) or PACAP (10 nM) for 3 h prior to lysis and luminometry. Luciferase activities were normalized to the treatment response of the control. *B* and *C*, BDNF responses were enhanced by knockdown of KLHL15 and overexpression (OE) of B*’β* but diminished by KLHL15 overexpression or B*’β* knockdown. *D* and *E*, PACAP-stimulated ERK activities were modulated by KLHL15 but insensitive to B*’β* manipulation. Data points are values from individual wells of four independent experiments; also shown are means ± 95% CI. Data of the stimulated (+) conditions were analyzed by Kruskal-Wallis tests with Dunn’s multiple comparisons tests. \*\**p* < 0.01, \*\*\**p* < 0.001, and \*\*\*\**p* < 0.0001.

### A sequence and structure search identifies DCX and DCLK1/2 as potential KLHL15 substrates

To uncover additional neuronal KLHL15 substrates, we searched the human UniProt database for proteins that contain the FRY sequence, which is necessary for the interaction of KLHL15 with its two previously identified targets, PP2A/B*’β* (15) and the DNA endonuclease CtIP/RBBP8 (23). This search identified a list of 616 proteins, which was further narrowed down according to secondary structure prediction (Fig. 2A). The FRY sequence in B*’β* and CtIP/RBBP8 is predicted to adopt a *β*-strand flanked by extended unstructured regions, which we reasoned provides flexibility for protein– protein interactions. Constraining the list of FRY-containing proteins to those in which the tripeptide is within a 2-4 residue long *β*-strand surrounded by at least 5 unstructured residues on both sides yielded a list of 24 proteins (Fig. 2A), which includes signaling proteins, transcription factors, and proteins involved in proteostasis (Table 1). We focused on three proteins: DCX and two doublecortin-like kinases DCLK1, DCLK2, which belong to a group of 11 neuronal MAPs characterized by microtubule-binding, tandem DCX domains that function during brain development (25,33-37,39). The FRY tripeptide is situated at the extreme C-terminal border of the second DCX domain (Fig. 2B). This region is highly conserved with the FRY sequence present in all vertebrate orthologs of DCX, DCLK1, and DCLK2. The region is not conserved in DCLK3, an atypical and largely uncharacterized member of the DCX family, whose homology to DCLK1 and DCLK2 is largely confined to the kinase domain (40).

**Table 1.** List of 24 putative KLHL15 substrates identified by the two-step *in silico* screen.

| **Protein** | **Annotation** |
| --- | --- |
| ATXN2 | RNA stability regulator Ataxin-2/SCA2 |
| AXIN1 | β-catenin destruction complex component Axin-1 |
| CACNG8 | Voltage-dependent calcium channel γ8 subunit |
| CADH3 | Cadherin-3 |
| DCLK1 | Serine/threonine-protein kinase DCLK1 |
| DCLK2 | Serine/threonine-protein kinase DCLK2 |
| DCX | Neuronal migration protein doublecortin |
| DIXDC1 | Wnt signaling pathway effector Dixin |
| DUSP2 | Dual specificity protein phosphatase 2 |
| EPB41L2 | Erythrocyte membrane protein band 4.1 like 2 |
| HERC4 | Probable E3 ubiquitin-protein ligase HERC4 |
| PDE8 | Cyclic nucleotide phosphodiesterase 8B |
| PDZD8 | PDZ domain-containing protein 8 |
| PPP2R5B | Protein phosphatase 2A regulatory subunit B'$\beta$ |
| PPIAL4C | Peptidyl-prolyl cis-trans isomerase A-like 4C |
| PSMG2 | Proteasome assembly chaperone 2 |
| PTPN4 | Tyrosine-protein phosphatase non-receptor type 4 |
| RABL6A | RAB-like GTPase RABL6A |
| RBBP8 | DNA endonuclease RBBP8/CtIP |
| TCF20 | Transcription factor 20 |
| TMEM161A | Transmembrane protein 161A |
| TRPS1 | Zinc finger transcription factor Trps1 |
| USPL1 | SUMO-specific isopeptidase USPL1 |
| ZFP37 | Zinc finger protein 37 |

**Figure 2.**
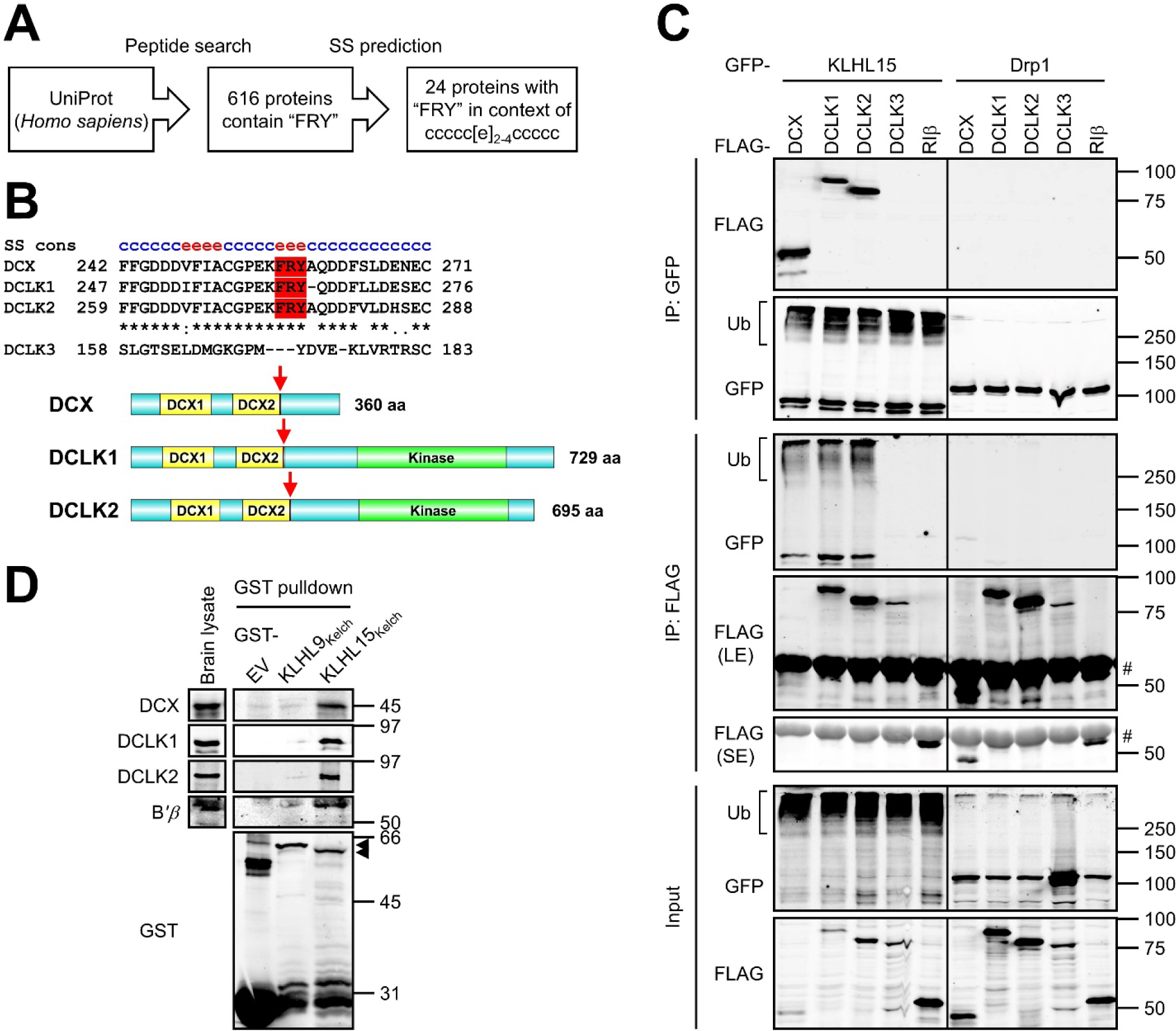
KLHL15 interacts with DCX, DCLK1, and DCLK2. *A*, search strategy for putative KLHL15 substrates. The human UniProt database was first screened for proteins containing the FRY sequence. The resulting 616 proteins were analyzed by consensus secondary structure (SS) prediction (https://npsa-prabi.ibcp.fr/) and filtered for proteins in which the FRY sequence is centered in the listed secondary structure (c = random coil, e = *β*-strand), yielding the 24 proteins listed in Table 1. *B*, on top is a multiple sequence alignment of human DCX (UniProt ID: O43602-2), DCLK1 (UniProt ID: O15075-2), DCLK2 (UniProt ID: G5E9L9) with FRY sequence in red and consensus secondary structure (cons SS) above. Because of insufficient homology, DCLK3 (UniProt ID: Q9C098) was aligned manually. Domain diagrams with red arrows indicating the FRY sequence are shown at the bottom. *C*, GFP-tagged KLHL15 was coexpressed with FLAG-tagged DCX, DCLK1, DCLK2, or DCLK3 in COS-1 cells, and KLHL15–substrate interaction was assessed by reciprocal immunoprecipitation (IP) via GFP or FLAG epitope tag and immunoblotting with the indicated antibodies. GFP-Drp1 and FLAG-RI*β* were included as a negative control for GFP or FLAG protein, respectively. Western blots representative of four independent experiments are shown. Ub: polyubiquitinated species; LE: long exposure; SE: short exposure; #: IgG heavy chain. *D*, bacterially expressed GST, GST-KLHL9_Kelch_, or GST-KLHL15_Kelch_ protein was incubated with a whole brain lysate from E18 rat embryo, and association of endogenous proteins was evaluated by GST pulldown and immunoblotting with the indicated antibodies. Western blots representative of three independent experiments are shown. Arrows point to the full-length GST-KLHL9_Kelch_ and GST-KLHL15_Kelch_ proteins. Molecular mass marker positions are indicated in kDa.

### KLHL15 interacts with DCX, DCLK1, and DCLK2

To test whether DCX, DCLK1, and DCLK2 are KLHL15 interactors, we co-expressed GFP-tagged KLHL15 and FLAG-tagged DCX or DCLKs (human cDNAs) in COS-1 cells and conducted reciprocal co-immunoprecipitation (co-IP) experiments. GFP-IP of KLHL15 selectively enriched DCX, DCLK1, and DCLK2, but not DCLK3 or another negative control (FLAG-RI*β*). Similarly, only FLAG-IPs of DCX and DCLK1/2 isolated GFP-KLHL15 (Fig. 2C). No interactions were detected with dynamin-related protein 1 (GFP-Drp1), indicating that DCX and DCLK1/2 do not interact with GFP. Notably, compared to GFP-Drp1, co-expression of GFP-KLHL15 dramatically decreased steady-state levels of the three interacting proteins (Fig. 2C, input).

To obtain evidence for direct interactions with endogenous proteins, we performed GST pulldown experiments with brain lysates from rat embryos (E18). A GST fusion of the substrate-binding Kelch domain of KLHL15 (amino acids 255–604, GST-KLHL15_Kelch_ (15)) was able to capture the endogenous PP2A/B*’β* regulatory subunit, as well as DCX, DCLK1, and DCLK2 (Fig. 2D). Indicating specificity for KLHL15, none of the proteins were pulled down with the Kelch domain of KLHL9 (GST-KLHL9_Kelch_) or GST alone (EV, Fig. 2D).

### The tandem DCX domains are necessary and sufficient for KLHL15 binding

In order to map the KLHL15-binding domain of DCX proteins, we constructed chimeras between the interactor DCLK2 and the non-interactor DCLK3. Chimera DCLK3-2 is composed of the N-terminus of DCLK3 (amino acids 1–130) and C-terminus of DCLK2 (amino acids 236–695), including the FRY tripeptide and the C-terminal half of the second DCX domain (Fig. 3A). DCLK2-3 is a chimera between the DCLK2 N-terminus (amino acids 1–281; including DCX1 and DCX2) and the C-terminal Ser/Thr kinase domain of DCLK3 (amino acids 176–648, Fig. 3A). According to co-IP experiments with transfected COS-1 cells, GFP-KLHL15 immunoisolated comparable levels of DCLK2 and DCLK2-3 (Fig. 3B, labeled FL = full-length). In contrast, DCLK3-2 was not detectable in the GFP-KLHL15 IP. These data indicate that DCLK2_1-281_ mediates the interaction with KLHL15, but that residues 236–281 including the FRY sequence are not sufficient for binding.

**Figure 3.**
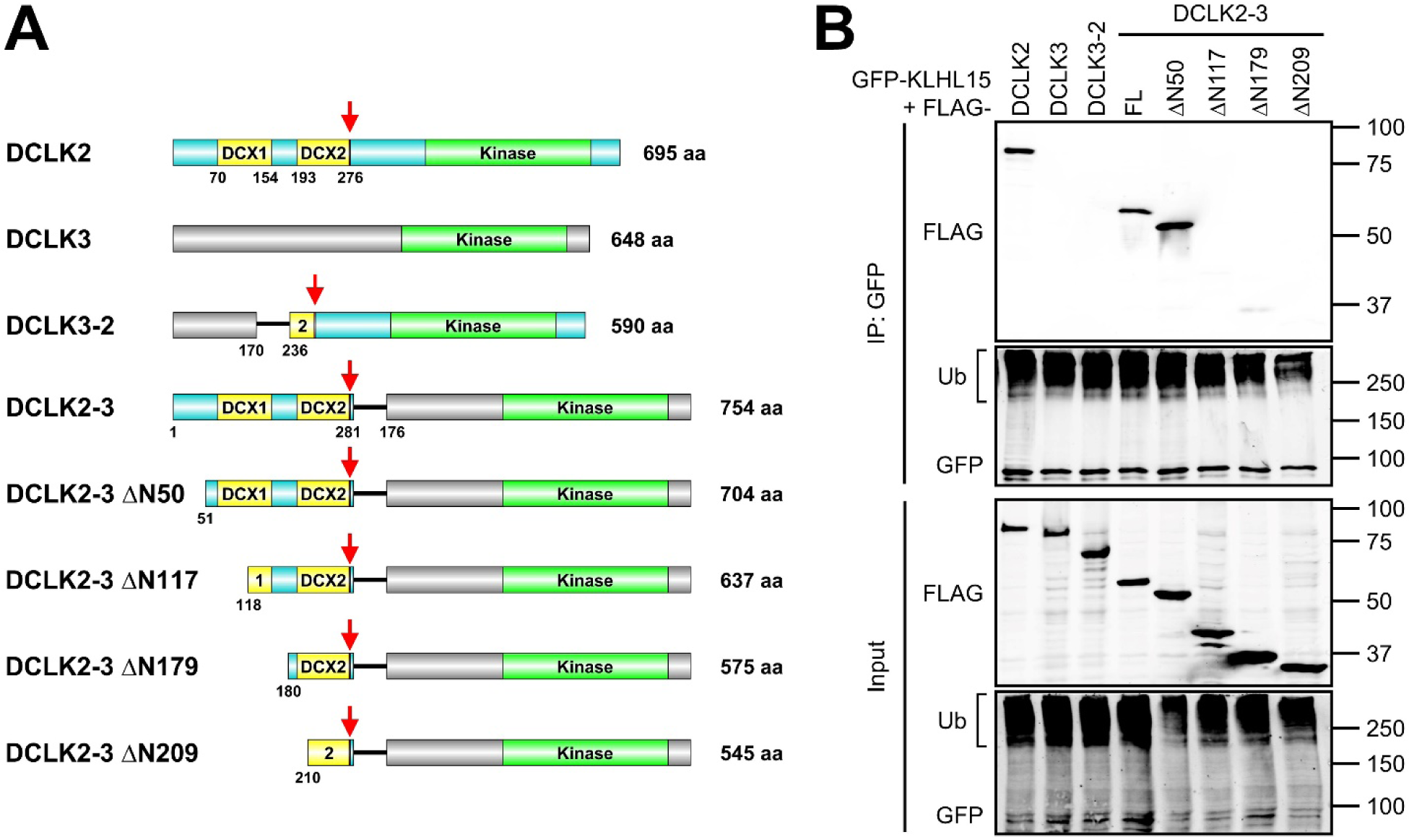
The minimal KLHL15 docking region of DCLK2 includes both DCX domains. *A*, domain diagrams of DCLK2, DCLK3, and full-length or truncated DCLK chimeras. Red arrows mark the locations of the FRY sequence. *B*, GFP-tagged KLHL15 was coexpressed with the indicated FLAG-tagged DCLK constructs in COS-1 cells, and interactions were analyzed by GFP-IP and probing for FLAG. Shown are Western blots representative of four independent experiments. Ub: polyubiquitinated species; FL: full-length. Molecular mass marker positions are shown in kDa.

To further narrow down the KLHL15 docking region within the DCLK2 N-terminus, we generated N-terminal truncations in the context of the DCLK2-3 chimeric cDNA (Fig. 3A). Deleting 50 amino acids from the unstructured N-terminus did not compromise the association with GFP-KLHL15 (Fig. 3B, compare lanes FL and ΔN50). Truncations into the first DCX domain (ΔN117) and beyond (ΔN179, ΔN209), in contrast, eliminated KLHL15 binding (Fig. 3B). Thus, DCLK2 residues 51–281 including both DCX domains are necessary and sufficient for the KLHL15 interaction. These mapping results likely also apply to DCX and DCLK1, in which this region is 80% and 82% identical, respectively.

### Tyr within the FRY sequence of DCX proteins is critical for KLHL15-mediated poly-ubiquitination and degradation

Point mutations within the FRY sequences of B*’β* and CtIP/RBBP8 disrupt interactions with KLHL15 (15,23). To evaluate whether this also holds true for DCX proteins, we generated Y to L substitutions within their FRY sequences. Indeed, DCX-Y259L, DCLK1-Y265L, and DCLK2-Y276L displayed significantly decreased KLHL15 binding in co-IPs (∼10-20% of their wild-type counterparts, Fig. 4, A and B), highlighting the critical importance of the Tyr residue in the FRY sequence. Residual KLHL15 binding of the mutant proteins is likely a consequence of their increased steady-state levels (Fig. 4A, input).

**Figure 4.**
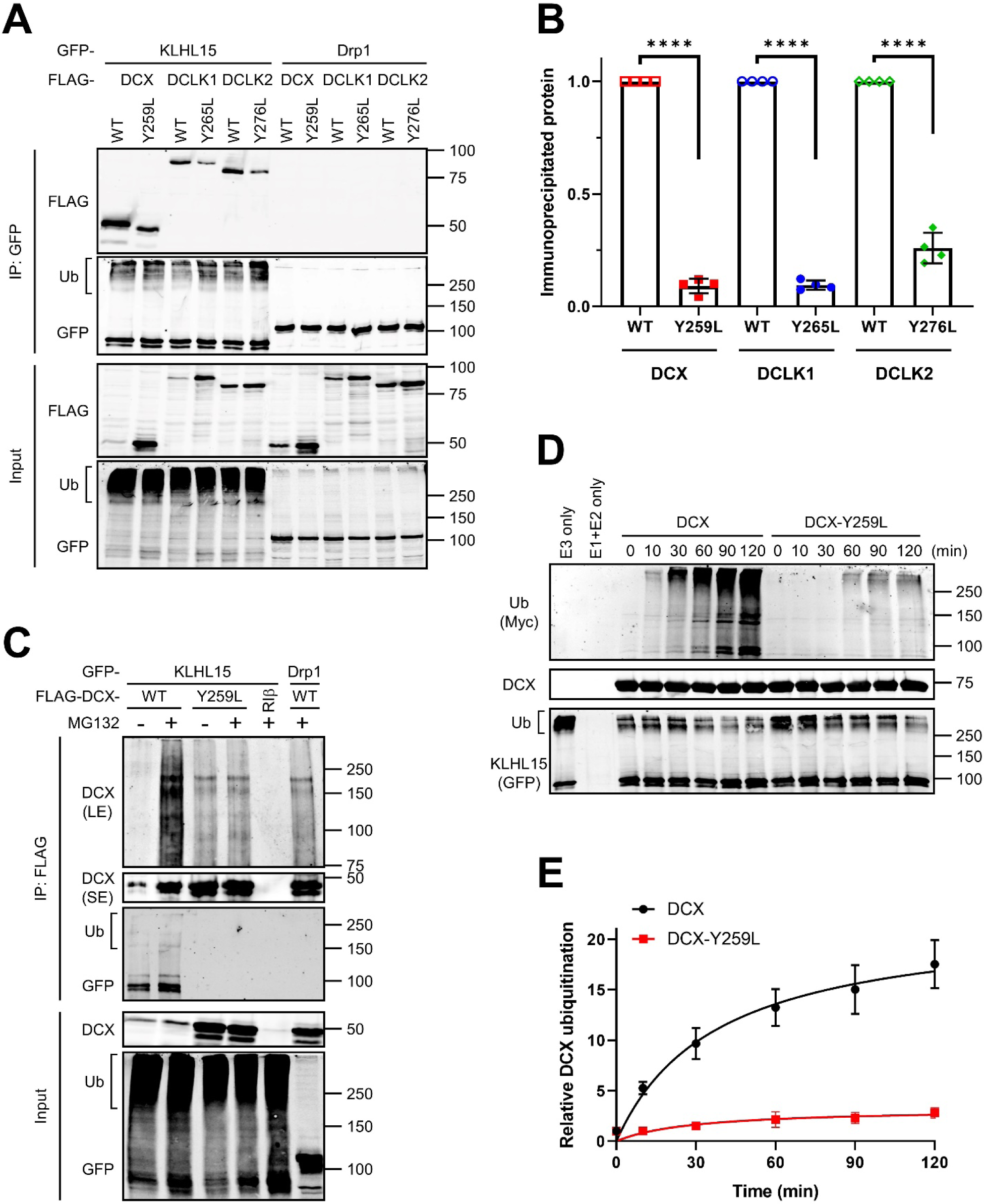
The FRY sequence of DCX is critical for KLHL15 binding and KLHL15-mediated polyubiquitination in cells and in *in vitro*. *A*, GFP-tagged KLHL15 or Drp1 was coexpressed with FLAG-tagged DCX, DCLK1, or DCLK2, either wild-type (WT) or FRY-mutant in COS-1 cells, and interaction was assessed by GFP-IP and immunoblotting with the indicated antibodies. *B*, relative DCX (DCLK) level after IP was quantified by dividing IP’d FLAG signals to IP’d GFP signals and then to FLAG signals in the corresponding input lane (FLAG^IP^/GFP^IP^/FLAG^Input^) and normalizing to the WT group. Shown are means ± SD as well as individual data points from four independent experiments. \*\*\*\**p* < 0.0001 by Student t-test. *C*, COS-1 cells coexpressing GFP-tagged KLHL15 and FLAG-tagged DCX WT or Y259L were treated with and without the proteasome inhibitor MG132 and lysates were subjected to FLAG-IP followed by immunoblotting with the indicated antibodies. GFP-Drp1 and FLAG-RI*β* were included as a negative control for GFP or FLAG protein, respectively. The unmodified DCX was detected by a short exposure (SE), whereas polyubiquitinated DCX was visualized by long exposure (LE). Shown are Western blots representative of four independent experiments. *D*, Myc-tagged ubiquitin and purified E1/E2 enzymes were mixed with immunoisolated Cul3– KLHL15 E3 ubiquitin ligase and bacterially expressed DCX-HaloTag-His_6_ protein (WT or Y259L), and *in vitro* ubiquitination reactions (37 °C for the indicated times) was initiated by addition of ATP. Molecular mass marker positions are shown in kDa. *E*, the time course of *in vitro* DCX ubiquitination was quantified by dividing Myc signals to DCX signals and GFP signals in the same lane (Myc/DCX/GFP) and normalizing to the zero time point (set as 1). Data are shown as means ± SD from four independent experiments and were fitted to the Michaelis– Menten curve using GraphPad Prism.

We next tested whether KLHL15 binding mediates polyubiquitination of the DCX proteins, first in an intact-cell context. COS-1 cells were cotransfected with HA-tagged ubiquitin, GFP-tagged KLHL15, and FLAG-tagged wild-type or FRY-mutant DCX and treated with proteasome inhibitor MG132 to allow accumulation of ubiquitinated proteins. Polyubiquitinated species of DCX were concentrated by FLAG-IP and probed with a DCX-specific antibody. Compared to GFP-Drp1, overexpression of KLHL15 promoted robust DCX ubiquitination (Fig. 4C). Polyubiquitination was substantially reduced by the Y259L mutation, despite higher levels of the unmodified, mutant protein in the IP (Fig. 4C), again underscoring the importance of the FRY sequence for substrate recognition and ubiquitin transfer by KLHL15.

Next, we inquired whether the Cul3– KLHL15 E3 ubiquitin ligase can catalyze DCX ubiquitination *in vitro*. To this end, the ubiquitination reaction was reconstituted by combining Myc-tagged ubiquitin, E1 ubiquitin-activating enzyme, E2 ubiquitin-conjugating enzyme, and immunopurified Cul3–KLHL15 E3 ubiquitin ligase with bacterially-expressed DCX-HaloTag fusion protein (wild-type or Y259L mutant), and initiated by the addition of ATP. Incubation of Cul3–KLHL15 E3 ligase with the wild-type DCX protein induced robust and time-dependent DCX polyubiquitination. In contrast, ubiquitination of DCX Y259L was strongly impaired under these conditions (Fig. 4, D and E), demonstrating that the Cul3– KLHL15 E3 ubiquitin ligase catalyzes DCX ubiquitination in a Y259-dependent manner.

To investigate DCX protein turnover and its dependence on KLHL15 and the FRY sequence, we analyzed substrate protein degradation using a HaloTag-based pulse-chase protocol, which we recently developed for long-lived proteins (41). HEK293T cells expressing endogenous KLHL15 (15) were cotransfected with DCX-HaloTag fusion cDNAs (wild-type or Y259L) and either GFP-KLHL15 or KLHL15-directed shRNAs (15), followed by pulse-labeling with a cell-permeant and covalent HaloTag ligand coupled to the fluorophore tetramethyl rhodamine (TMR). Labeled DCX-HaloTag was chased by incubating cells with the non-fluorescent competitive ligand 7-bromoheptanol for up to 24 h (41). DCX turnover was quantified by dividing TMR fluorescence by DCX immunoblot signals in the same lane and normalizing to the zero time point. Silencing endogenous KLHL15 with two different shRNAs increased steady-state levels of wild-type DCX-HaloTag (TMR labeling without chase) and markedly slowed its turnover (*t*_1/2_ > 24 h versus *t*_1/2_ = 16 h, from monoexponential decay curve fits). In contrast, KLHL15 silencing did not further stabilize the already long-lived Y259L-mutant DCX (all *t*_1/2_ > 24 h; Fig. 5, A and B). Conversely, overexpression of GFP-KLHL15 lowered wild-type DCX steady-state levels and accelerated DCX turnover (*t*_1/2_ = 5 h; Fig. 5, A and B), while having minimal effects on turnover of the FRY-mutant protein (*t*_1/2_ = 20 h; Fig. 5, A and B). Statistical analyses of areas under the degradation curves (AUCs, Fig. 5C) confirmed our identification of KLHL15 as a critical determinant of DCX stability and the importance of the FRY tripeptide.

**Figure 5.**
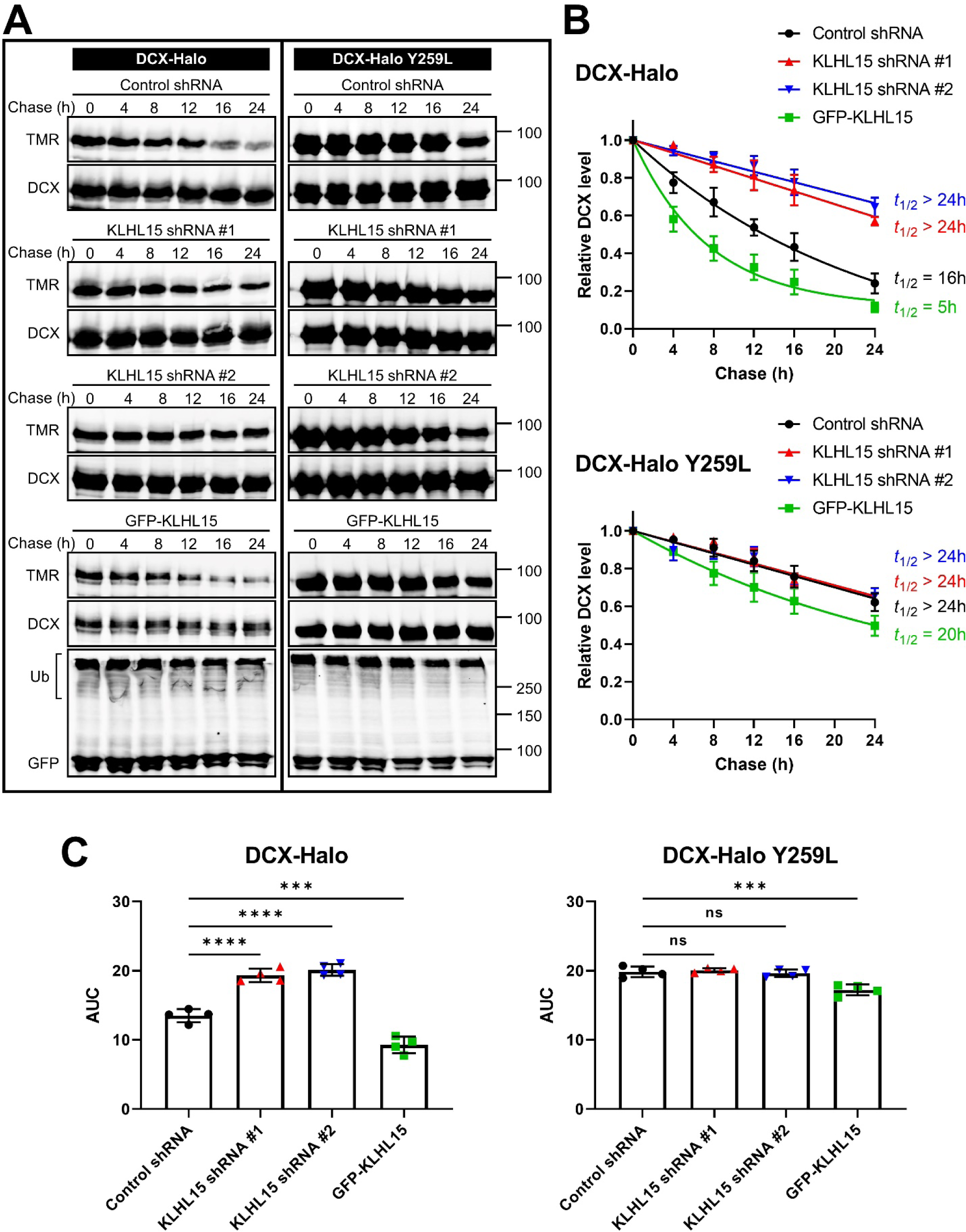
KLHL15 promotes DCX protein turnover in a FRY-dependent manner. HEK293T cells were cotransfected with DCX-HaloTag (WT or Y259L) and either the indicated shRNA plasmids or GFP-KLHL15. DCX-HaloTag was pulse-labeled (15 min, 100 nM) with tetramethyl rhodamine (TMR)-labeled HaloTag ligand and then chased with the non-fluorescence ligand 7-bromoheptanol for the indicated times before cells were lysed in SDS. *A*, total lysate blots of cells expressing WT (left) or mutant (right) DCX-HaloTag were first imaged for TMR fluorescence and then immunoblotted for DCX and GFP. Molecular mass marker positions are shown in kDa. *B*, DCX degradation was quantified by dividing TMR fluorescence by DCX immunoblotting signals in the same lane (TMR/DCX) and normalizing to the zero time point. Data points are means ± SD from four independent transfections; half-lives (*t*_1/2_) were determined from the shown one-phase decay curve fits. *C*, area-under-the-curve (AUC) was quantified from the decay curves in *B* and is plotted as individual data points and means ± SD from four independent experiments. *p* values were obtained by one-way ANOVA followed by Dunnett’s post-hoc tests for multiple comparisons to the control shRNA group. \*\*\**p* < 0.001, \*\*\*\**p* < 0.0001.

We similarly examined turnover of the two DCX-domain containing protein kinases using HaloTag pulse-chase labeling in HEK293T cells. With half-lives of greater than 24 h, both DCLK1 (Fig. 6) and DCLK2 (Fig. 7) were significantly more stable than DCX (greater AUCs, *p* < 0.0001). Nonetheless, forced expression of GFP-KLHL15 promoted degradation of both DCLK1 (*t*_1/2_ = 10 h) and DCLK2 (*t*_1/2_ = 18 h), whereas endogenous KLHL15 silencing further stabilized the DCX-domain kinases as evidenced by AUC analyses (Fig. 6C, 7C). Just like DCX-Y259L, the KLHL15-binding impaired DCLK1-Y265L and DCLK2-Y276L mutants were not further stabilized by KLHL15 silencing. As well, turnover of mutant DCLK1/2 was less sensitive to KLHL15 overexpression than turnover of their wild-type counterparts (AUCs, *p* < 0.01). In aggregate, these data indicate that protein stability of DCX, DCLK1, and DCLK2 is controlled by KLHL15 in a FRY-dependent manner.

**Figure 6.**
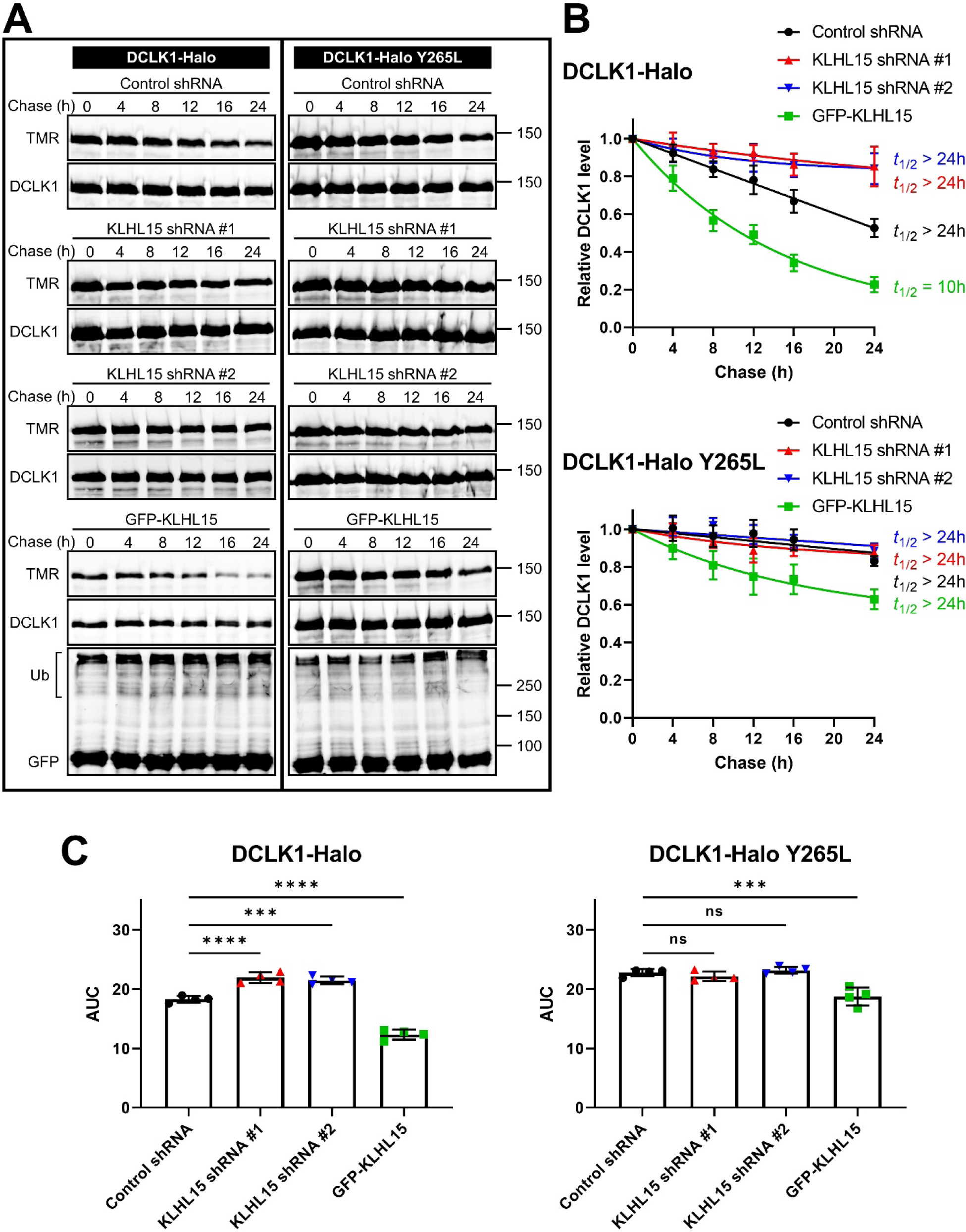
KLHL15 accelerates DCLK1 degradation in a Y265-dependent manner. HEK293T cells were cotransfected with DCLK1-HaloTag (WT or Y265L) and either GFP-KLHL15 or the indicated shRNA plasmids. DCLK1-HaloTag was pulse-labeled (15 min, 100 nM) with TMR-labeled HaloTag ligand and then chased with the 7-bromoheptanol for the indicated times before cell lysis. *A*, total lysate blots of cells expressing WT (left) or mutant (right) DCLK1-HaloTag were first imaged for TMR fluorescence and then immunoblotted for DCLK1 and GFP. Molecular mass marker positions are shown in kDa. *B*, DCLK1 degradation was quantified by dividing TMR fluorescence by DCLK1 immunoblotting signals in the same lane (TMR/DCLK1) and normalizing to the zero time point. Data points are means ± SD from four independent transfections; half-lives (*t*_1/2_) were determined from the shown one-phase decay curve fits. *C*, area-under-the-curve (AUC) was quantified from the decay curves in *B* and is plotted as individual data points and means ± SD from four independent experiments. Data were analyzed by one-way ANOVA followed by Dunnett’s post-hoc tests for multiple comparisons to the control shRNA group. \*\*\**p* < 0.001, \*\*\*\**p* < 0.0001.

**Figure 7.**
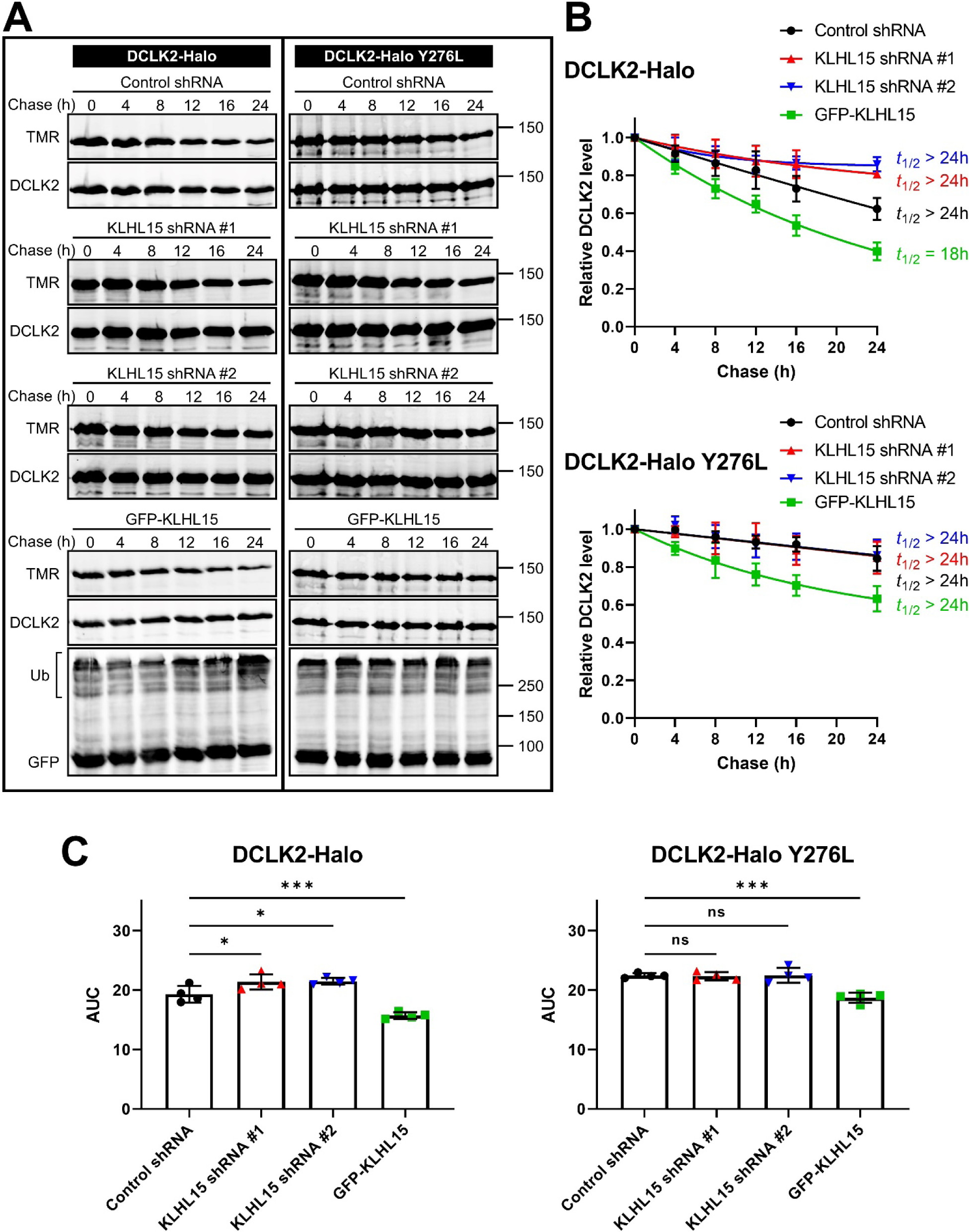
KLHL15 promotes DCLK2 protein turnover in a Y276-dependent manner. HEK293T cells were cotransfected with DCLK2-HaloTag (WT or Y276L) and either the indicated shRNA plasmids or GFP-KLHL15. DCLK2-HaloTag was pulse-labeled (15 min, 100 nM) with TMR-labeled HaloTag ligand and then chased with the non-fluorescence ligand 7-bromoheptanol for the indicated times before cell lysis. *A*, total lysate blots of cells expressing WT (left) or mutant (right) DCLK2-HaloTag were first imaged for TMR fluorescence and then immunoblotted for DCLK2 and GFP. Molecular mass marker positions are shown in kDa. *B*, DCLK2 degradation was quantified by dividing TMR fluorescence by DCLK2 immunoblotting signals in the same lane (TMR/DCLK2) and normalizing to the zero time point. Data points are means ± SD from four independent transfections; half-lives (*t*_1/2_) were determined from the shown one-phase decay curve fits. *C*, area-under-the-curve (AUC) was quantified from the decay curves in *B* and is plotted as individual data points and means ± SD from four repeated experiments. Data were analyzed by one-way ANOVA followed by Dunnett’s post-hoc tests for multiple comparisons to the control shRNA group. \*\*\**p* < 0.001, \*\*\*\**p* < 0.0001.

### KLHL15 negatively regulates dendritic complexity of hippocampal neurons

DCX is an abundant microtubule-binding protein in the developing brain, whose loss of function leads to severe neurological deficits in humans (28,42). To examine the interplay between the two XLID genes *DCX* and *KLHL15* in a model of neuronal development, we quantified dendritic complexity of primary hippocampal neurons cultured from E18 rat embryos. Two days after plating, cultures were transfected with membrane-targeted mCherry (Lck-mCherry) to visualize neurites, together with GFP-KLHL15 and either wild-type or FRY-mutant DCX, or empty vector (GFP, EV) controls. Three days later, neurons were fixed, imaged, and dendritic trees were traced semi-automatically for total length and Sholl analysis. The expression of either wild-type or mutant DCX alone resulted in a slight, but non-significant increase in total dendritic intersections and total dendrite length compared to the control (Fig. 8, B–E), likely due to a ceiling effect of high endogenous DCX levels in these embryonic cultures. However, coexpression of GFP-KLHL15 with wild-type DCX-WT dramatically reduced dendritic complexity and length. Dendritic growth stunted by KLHL15 could be completely reversed by coexpressing the KLHL15-resistant DCX-Y259L mutant protein. These results suggest that KLHL15 restricts process formation in embryonic neurons by targeting DCX for proteasomal degradation.

**Figure 8.**
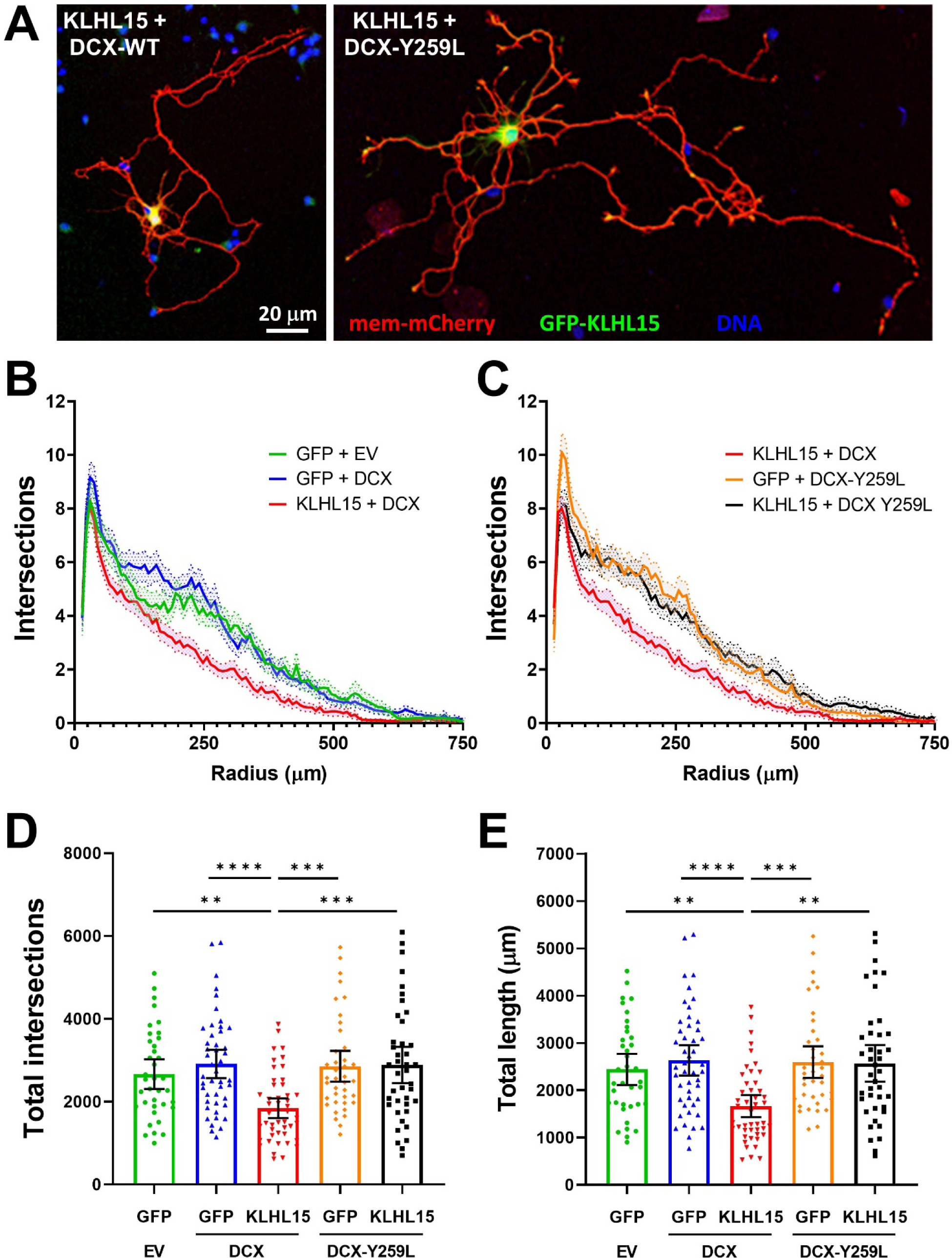
KLHL15 antagonizes dendrite outgrowth of hippocampal neurons via DCX. *A*, representative images of hippocampal neurons from E18 rat embryos transfected with membrane-targeted mCherry (red), GFP-KLHL15 (green) and either DCX-WT (left) or DCX-Y259L (right); nuclei are labeled blue. *B* and *C*, plots of numbers of dendrite intersections with concentric circles (steps of 7.5 μm starting 15 μm from the soma) obtained by Sholl analysis of traced dendrites. For clarity, the five transfection conditions were separated into two graphs with KLHL15 + DCX shown in both. *D* and *E*, plots of the total number of dendrite intersections from the Sholl analysis (D) and total dendrite length (E). Shown are means ± 95% CI (as well as individual data points in D and E) of 37–47 neurons from three or four cultures per transfection condition. Data were analyzed by Kruskal-Wallis followed by Dunn’s multiple comparisons tests. \*\**p* < 0.01, \*\*\**p* < 0.001, and \*\*\*\**p* < 0.0001.

## DISCUSSION

Ubiquitin-processing enzymes are fundamental for neuronal development and function (43,44) and dysregulation of these enzymes can cause neurodevelopmental (2,3,45) and neurodegenerative diseases (4,46). E3 ligases determine the substrate specificity of the ubiquitination reaction, as they directly interact with proteins targeted for ubiquitination (47). Among >600 predicted human E3 ligases, Cullin-RING ligases (CRLs) constitute the largest family, regulating a plethora of biological pathways (48,49). In fact, mutations in many CRL genes are strongly associated with neurodevelopmental disorders (5-7,50).

Among the cullin3-based CRLs, E3 ubiquitin ligases of the KLHL (BTB-Kelch) family catalyze the formation of predominantly lysine 48-linked ubiquitin chains that trigger proteasomal degradation (12-14). Many *KLHL* genes have been implicated in carcinogenesis and hereditary disorders (12) and loss-of-function of *KLHL15*, the subject of the present study, is associated with severe XLID (8).

Here, we used an *in silico* approach to identify novel substrates of KLHL15 important for early brain development. Our results show that KLHL15 interacts with and targets three of the eleven members of the doublecortin family of neuronal MAPs (DCX, DCLK1, and DCLK2) for ubiquitination and subsequent proteasomal degradation. Identified now in a total of five KLHL15 targets, including the PP2A regulatory subunit B*’β* and the DNA repair protein CtIP/RBBP8 (15,23), a FRY tripeptide embedded within a predicted unstructured region is required for recognition of the three DCX domain-containing proteins by the E3 ligase.

The KLHL15 binding domain, or degron, we identified in DCX (residues 51 to 281) encompasses the FRY sequence at its C-terminal end, but also the protein-family defining, tandem DCX domains, which have been shown to stabilize the microtubule cytoskeleton by binding selectively to 13 protofilament-containing microtubules (51-53). The overlap between microtubule- and KLHL15-binding domains of DCX suggests the possibility of competitive interactions, for instance, KLHL15 may only ubiquitinate DCX proteins that are not bound to microtubules.

DCX is developmentally regulated in the mammalian brain with high expression in embryonic stages followed by sharp decline in adulthood (54-59). DCX is also known to have heterogenous subcellular distributions in various neuronal cell types. DCX is particularly enriched at the ends of neuronal processes where microtubules enter the growth cone (60) and axonal segments that can generate collaterals (61). Moreover, DCX is further involved in nuclear translocation during neuroblast migration and may influence neuroblast differentiation (62). Distinct patterns of expression and localization strongly implicate the spatiotemporal regulation of DCX via the ubiquitin-proteasome pathway. DCX’s two paralogs, DCLK1 and DCLK2 also display developmental stage-specific expression and preferentially localize to distal dendrites to promote their growth by enhancing microtubule bundling (33), implying similar proteolytic regulation of the DCX domain-containing kinases.

DCX domain proteins are likely targeted by multiple E3 ligases that are themselves differentially expressed and localized. Indeed, the nuclear E3 ubiquitin ligase Mdm2 has been reported to ubiquitinate and degrade DCX in newborn olfactory bulb interneurons (63). Mechanisms for DCX stabilization have also been documented. For instance, the 14-3-3ε protein binds and protects DCX from ubiquitination and subsequent degradation (64), while the deubiquitinating enzyme USP9X binds DCX in a ubiquitination-independent manner (57).

Reporter assays in hippocampal neurons point to KLHL15 as a potent negative regulator of BDNF/TrkB and PACAP/PAC1 signaling to ERK (Fig. 1). Knockdown and overexpression of the KLHL15 target B*’β* suggests that KLHL15-mediated inhibition of TrkB signaling likely occurs via degradation of PP2A/B*’β*, a neuronal protein phosphatase which we previously reported amplifies NGF/TrkA signaling in PC12 cells (22). On the other hand, PACAP-mediated ERK stimulation is inhibited by KLHL15 independently of this PP2A regulatory subunit. While DCX-domain kinases have previously been reported to modulate transcriptional response (40), it is unclear whether DCX proteins or other, as yet to be identified substrates of the E3 ligase mediate KLHL15’s effects on PACAP signaling.

DCX plays pivotal roles in neurogenesis, neuronal migration, and process outgrowth to establish cortical and hippocampal lamination during early brain development (28,29). In particular, DCX knockout or RNAi-mediated silencing impairs neurite outgrowth in developing cortical and hippocampal neurons (42,65). Consistent with DCX stabilizing dendritic microtubules and KLHL15 targeting DCX for proteasomal degradation, we found that overexpression of KLHL15 reduces dendritic complexity in hippocampal neurons expressing wild-type, but not FRY-mutant DCX (Fig. 6).

Whereas *DCX* inactivating mutations cause severe structural brain abnormalities (30), the case for *DCLK1* and *DCLK2* as disease genes for neurodevelopmental disorders is currently circumstantial (66,67). While several studies identified *KLHL15* as an X-linked gene associated with various neurodevelopmental complications (8-11), a connection to DCX domain proteins is not intuitively obvious, as their levels would be expected to increase, rather than decrease in the absence of the E3 ligase. One could envision scenarios, however, in which *KLHL15* inactivation interferes with the normal developmental downregulation of DCX proteins and their replacement on microtubules with MAPs expressed later in development. Further studies involving animal models are required to investigate the interplay between KLHL15 and DCX proteins in the pathogenesis of neurodevelopmental disorders.

## EXPERIMENTAL PROCEDURES

### Cell culture and transfection

COS-1 and HEK293T cells were cultured at 37 °C and 5% CO_2_ in Dulbecco’s Modified Eagle Medium (DMEM, Gibco) supplemented with 10% fetal bovine serum (FBS, Atlanta Biologicals) and 1% (v/v) GlutaMax (Gibco). Cells were grown to 60% confluency on collagen-coated plates and transfected using Lipofectamine 2000 (Invitrogen) following the manufacture’s protocol for transient transfection of adhered cells. Primary hippocampal cultures were prepared from embryonic day 18 (E18) Sprague Dawley rats (Harlan) as described previously (41). Hippocampi were dissected and pooled in ice cold HEPES-buffered neurobasal and then incubated in HBSS containing trypsin (1.5 mg/ml) at 37 °C for 12 min. The tissues were washed three times with HBSS before cells were dissociated by trituration. Cells were plated in Neurobasal complete (Neurobasal media (Gibco), 2% B27 supplement, 10 mM HEPES, 0.25% GlutaMax), and 5% horse serum on poly-l-lysine-coated plates. Medium was changed to serum-free Neurobasal complete after 4 h of incubation. Cells were maintained at 37 °C in a humidified environment of 95% air/5% CO_2_ with half the media changed with fresh media twice a week (41).

### Plasmids

The green fluorescent protein (GFP)- tagged KLHL15 replacement plasmids, control, KLHL15- and B*’β*-directed small hairpin RNAs (shRNAs) were previously described (15). The human doublecortin (DCX) and doublecortin-like kinases (DCLK1, DCLK2, and DCLK3) cDNAs were obtained from Origene in pCMV6 vector that also incorporates C-terminal Myc and FLAG tags (Myc-FLAG) or subcloned into a pHTC (Promega)-derived vector that expresses C-terminal HaloTag (41). Missense mutations, chimeras, and truncations of DCX and DCLKs were generated by PCR following standard protocol (15). The DCX-HaloTag-His_6_ fusion cDNA was also subcloned into the bacterial expression plasmid pET-21b(+) using compatible restriction sites. Plasmids expressing GFP-Drp1, GFP-B*β*2 RR168EE, FLAG-RI*β*, GST-KLHL15_Kelch_, GST-KLHL9_Kelch_, HA-Cul3, HA-Cul3ΔRoc, and HA-ubiquitin were described previously (15). All constructs were verified by sequencing by the Iowa Institute of Human Genetics (IIHG) Genomics Division.

### Antibodies and reagents

The commercially-sourced primary antibodies used in this study include: rabbit anti-GFP (ab290, Abcam), mouse anti-GFP (N86/8) (75-131, NeuroMab), mouse anti-FLAG (M2) (F3165, Sigma), rabbit anti-DYKDDDDK (FLAG) Tag (2368, Cell Signaling), mouse anti-Doublecortin (E-6) (sc-271390, Santa Cruz), rabbit anti-Doublecortin (4604, Cell Signaling), rabbit anti-DCLK1/DCAMKL1 (D2U3L) (62257, Cell Signaling), rabbit anti-DCLK2 (ab106639, Abcam), goat anti-GST (27-4577-01, Amersham), mouse anti-V5-Tag (46-0705, Invitrogen), mouse anti-Myc-Tag (9B11) (2276, Cell Signaling), rabbit anti-HA-Tag (C29F4) (3724, Cell Signaling), and rabbit anti-ubiquitin (3933, Cell Signaling). The rabbit polyclonal antibody against B*’β* was described previously (16). Secondary antibodies were: IRDye® 680RD goat anti-mouse IgG (926-68070, LI-COR Biosciences), IRDye® 800CW goat anti-rabbit IgG (926-32211, LI-COR Biosciences), IRDye® 680RD donkey anti-mouse IgG (926-68072, LI-COR Biosciences), IRDye® 800CW donkey anti-rabbit IgG (926-32213, LI-COR Biosciences), IRDye® 680RD donkey anti-goat IgG (926-68074, LI-COR Biosciences), and Superclonal™ goat anti-rabbit IgG, HRP conjugate (A27036, Invitrogen).

The chemical reagents used in this study were listed as follows: MG132 (C2211, Sigma), MLN4924 (5.05477.0001, Millipore), HaloTag® TMR ligand (G8251, Promega), 7-Bromo-1-heptanol (H54762, Alfa Aesar), EZview™ Red anti-FLAG® M2 affinity gel (F2426, Sigma), GFP-nanobody agarose resin (143093, UI Biomedical Research Store), Glutathione-Superflow resin (635607, Takara), His60 Ni Superflow Resin (635660, Takara), lysozyme from chicken egg white (L6876, Sigma), recombinant human UBE1 (E-305, Boston Biochem), recombinant human UbcH5a/UBE2D1 (E2-616, Boston Biochem), Myc-ubiquitin (U-115, Boston Biochem), ubiquitin aldehyde (U-201, Boston Biochem), creatine phosphokinase from rabbit muscle (C3755, Sigma), sodium creatine phosphate dibasic tetrahydrate (27920, Sigma), ATP (R0441, Thermo Scientific), human BDNF (2837, Tocris), PACAP-38 (4031157, Bachem), poly-L-lysine hydrobromide (OKK-3056, Peptides international), and Western Lightning® Ultra (NEL113001EA, PerkinElmer).

### Dual luciferase reporter assays

Primary rat hippocampal cultures were plated at 50,000 cells/well in 48 well plates and transfected on DIV 9 (day in vitro) using NeuroMag magnetic beads. For the transfection, KLHL15, B*’β* or GFP and empty vector control or shRNA-expressing plasmid were combined with the PathDetect Elk1 trans-reporting system (Agilent) and a SV40-pNL1.1, for the normalization. On DIV13, cultures were treated with either media, or media containing BDNF (25 ng/ml final) or PACAP (10 nM final) and assayed 3 hours later with the Nano-Glo® Dual-Luciferase® Reporter System (Cat# N1630; Promega) on a Berthold Sirius luminometer to quantify ERK activation.

### Immunoprecipitation and immunoblot analyses

Immunoprecipitation (IP) was carried out as previously described (15). Briefly, COS-1 cells were seeded and grown on collagen-coated 6-well plates followed by transfection. Cells were lysed in IP buffer (20 mM Tris, pH 7.5, 150 mM NaCl, 1% Triton X-100, 1 mM EDTA, 1 mM EGTA, 1 mM benzamidine, 1 mM PMSF, and 1 μg/ml leupeptin) and rotated at 4 °C for 30 min to allow sufficient solubilization. The total lysates were centrifuged at 13,000 × *g* for 10 min to separate cytosol from insoluble fraction and cell debris. 10% of the supernatant was sampled out as input, the remainder was then subjected to either FLAG-IP or GFP-IP using pre-equilibrated anti-FLAG M2 affinity gel or GFP-nanobody agarose resin (10–15 μl bead volume per reaction), respectively, for 2 h rotating at 4 °C. Beads were washed 4 times by centrifugation in lysis buffer and eluted by boiling in 2× Laemmli sample buffer for 5 min. For *in vitro* ubiquitination assays, Cul3– KLHL15 E3 ubiquitin ligase complex was immunopurified by GFP-IP from HEK293T cells coexpressing GFP-KLHL15 and HA-Cul3. Precipitates were washed 4 times with 50 mM NaCl, 50 mM HEPES, 10% glycerol, 0.1% Tween 20, and 20 mM Tris, pH 7.5. The immobilized Cul3–KLHL15 E3 complex integrity was verified by immunoblotting and immediately used for *in vitro* ubiquitination assays. Samples were separated on 10% SDS-PAGE gels, electrotransferred to nitrocellulose membrane (GE Healthcare), and immunoblotted as indicated. Generally, proteins were visualized by using species-specific fluorescent secondary antibodies and a LI-COR Odyssey infrared scanner for dual-color detection. Immunoblotting signals were quantified by densitometry using the gel analysis plugin of ImageJ (National Institutes of Health).

### Purification of bacterially expressed proteins

GST, GST-KLHL9_Kelch_, GST-KLHL15_Kelch_, DCX-Halo-His_6_, DCX-Halo-His_6_ Y259L, and Halo-His_6_ were expressed in *E. coli* BL21(DE3) cells (200131, Agilent) and purified according to standard procedures as previously described (15). Briefly, bacterial cells were grown at 37 °C, 200 rpm to OD ∼0.8 and induced with 500 μM IPTG for 6–8 h.

Cells were pelleted by centrifugation at 3000 rpm for 15 min and lysed in 50 mM Tris, pH 7.5, 150 mM NaCl, 2 mM benzamidine, 1 mM PMSF, and 1 mg/ml lysozyme for 30 min to complete lysis, followed by a total of 2 min sonication (15 s on – 45 s off; 8 rounds on ice). Samples were centrifuged at 15,000 rpm for 15 min and supernatant was then applied to the appropriate resin following the manufacture’s protocol. The his-tagged proteins were eluted with 250 mM imidazole and dialyzed into the desired buffer while the GST proteins remained immobilized on glutathione resin. Purified proteins were quantified by Coomassie staining with BSA standards within the linear range and were utilized immediately.

### GST pulldown assays

Whole brains were rapidly dissected from embryos (E18) of Sprague-Dawley rats following decapitation, flash frozen in liquid nitrogen, and stored at -80 °C until use. For whole brain lysates, frozen tissues were pulverized and homogenized in lysis buffer containing 0.5% Triton X-100, 25 mM Tris, pH 7.5, 50 mM NaCl, 2 mM DTT, 0.5 mM EDTA, 1 mM PMSF, 1 mg/ml leupeptin, 1 mM benzamidine. Homogenates were sonicated to shear DNA and cleared by centrifugation (15 min at 20,000 × g). A portion of the brain homogenate supernatant was added directly to 4× Laemmli buffer to provide an input sample. GST pulldown assays were performed by mixing 60–100 μl whole brain lysates with ∼30 μg immobilized GST proteins for 2 h rotating at 4 °C. Glutathione resins were washed 4 times by centrifugation in lysis buffer and eluted in 2× Laemmli sample buffer. Protein samples were subjected to SDA-PAGE and immunoblotting with indicated antibodies.

### In-cell ubiquitination assays

24–36 h post-transfection, COS-1 cells were treated with 25 μM MG132 (from 50 mM stock in DMSO) for 8–12 h before lysis to permit sufficient proteasomal inhibition. A final concentration of 2 μM MLN4924 (from 10 mM stock in DMSO) was added alone or in combination with MG132 for NEDD8-activating enzyme (NAE) inhibition. Cytosolic fraction was then subject to FLAG-IP as described above and immunoblotted for DCX for DCX-specific ubiquitination. The effectiveness of MG132 was confirmed by the accumulation of DCX polyubiquitination and/or stabilization of steady-state protein level in the presence of MG132 in cells expressing FLAG-DCX alone. DCX immunoblotting signals were visualized by quantitative fluorescent imaging (LI-COR) using species-specific secondary antibodies to avoid detection of heavy chain and light chain of denatured antibodies during IP.

### In vitro ubiquitination assays

*In vitro* ubiquitination assays were done as previously described (68), but modified for 48-well plates. Briefly, the ubiquitination reaction mixture contained 125 ng of E1 enzyme (Boston Biochem), 250 ng of E2 enzyme (Boston Biochem), immunopurified Cul3–KLHL15 E3 complex, 10 μg of Myc-ubiquitin (Boston Biochem), 1 μM ubiquitin aldehyde (Boston Biochem), 10 mM MgCl_2_, 2 mM DTT, 10 mM creatine phosphate (Sigma), 0.5 mg/ml creatine phosphokinase (Sigma), and 20 μg of purified proteins including DCX-Halo-His_6_ (wild-type or Y259L) or Halo-His_6_ as control protein. The ubiquitination reactions were diluted to a final volume of 0.25 ml in 50 mM HEPES, pH 7.5, initiated upon addition of 5 mM ATP (Thermo Scientific), and incubated for up to 2 h at 37 °C on a titer plate shaker (constant speed 3, Lab-Line Instruments, Inc.). 30 μl reaction mixtures were sampled at the indicated times and immediately terminated in 4× Laemmli buffer. Proteins were then resolved on 8% gels by SDS-PAGE, and ubiquitinated proteins were detected by immunoblotting with Myc-Tag antibody.

Relative DCX ubiquitination *in vitro* was quantified by dividing Myc-Tag signals by DCX signals and GFP (KLHL15) signals in the same lane and normalizing to the zero time point (set to 1). The enzymatic kinetics was plotted in Michaelis–Menten model using GraphPad Prism.

### HaloTag turnover assays

HaloTag-based turnover assays were conducted following a pulse-chase protocol as previously reported (41). Briefly, HEK293T cells were transfected with HaloTag constructs with KLHL15-targeted shRNA or a control shRNA. For gain-of-function study, HaloTag construct was cotransfected with GFP-KLHL15 replacement vector or a GFP-control vector. 24–36 h post-transfection, cells were incubated with 50 nM fluorescent cell-permeant HaloTag® TMR ligand for 30 min at 37 °C to allow maximal labeling. Then TMR-containing media was removed and replaced with media containing 10 μM 7-Bromoheptanol (7-Br) blocker for the indicated times. The blocking was terminated by immediate cell lysis and HaloTag protein was separated by SDS-PAGE. TMR fluorescence was visualized by Sapphire Biomolecular Imager using a Cy3 filter (Azure Biosystems), and total DCX (or DCLK) protein was subsequently probed with indicated antibodies. Relative DCX (or DCLK) protein levels were quantified by dividing TMR fluorescence signals by individual immunoblotting signals in the same lane, and normalizing to the zero time point (set to 1). Half-lives (*t*_1/2_) were calculated from the one-phase decay model using GraphPad Prism.

### Dendrite outgrowth assays

Primary rat hippocampal cultures from E18 embryos were plates at 50,000 cells/ml in cover glass bottomed chambers and plates (C8-1.5H-N and P24-1.5H-N, Cellvis). At DIV 2, cultures were transfected with 0.2% Lipofectamine 2000 and the following DNA constructs at these ratios: 20% Lck-2V5-mCherry (plasma membrane targeted, also containing two V5 tags), 40% empty vector (EV), DCX-WT or DCX-Y259L, and 40% EGFP or GFP-KLHL15. On DIV 5, cultures were fixed in 4% PFA and intrinsic fluorescence was amplified with antibodies to GFP and the V5 tag and appropriate fluorescent secondary antibodies. Image stacks (11 focal planes at 1 μm intervals) were acquired using the 10X objective of a Keyence BZ-X800E microscope and converted to maximum intensity projections. Dendrites were traced semi-automatically using the ImageJ plugin SNT and traces were analyzed for length (total pixel number) and with the Sholl Analysis plugin for intersections with concentric circles (7.5 μm apart, starting 15 μm from the cell body). All analyses were performed blinded to the transfection conditions.

### Statistical analysis

Statistical analyses were performed using GraphPad Prism 8 with tests listed in the figure captions. The robust outlier removal (ROUT) method of removing outliers was used at Q = 1% prior to normalization of data and significance levels are abbreviated as follows: \**p* < 0.05, \*\**p* < 0.01, \*\*\**p* < 0.001, and \*\*\*\**p* < 0.0001.

## ACKNOWLEDGMENTS

This work was supported by National Institutes of Health grants DK116624, MH115673, MH113352 to S.S.. Additional support to S.S. was provided by Jordan’s Guardian Angels, the Roy J. Carver Charitable Trust, and Iowa Neuroscience Institute. We also acknowledge support by the Genomics Division of the UI Carver College of Medicine.

## AUTHOR CONTRIBUTIONS

Conceptualization: J.S., R.A.M., and S.S.; writing: J.S., R.A.M., and S.S.; investigation: J.S., R.A.M., and A.Y.U.; and resources; supervision; funding acquisition; project administration: S.S..

